# Large-scale genome-wide association meta-analysis of endometriosis reveals 13 novel loci and genetically-associated comorbidity with other pain conditions

**DOI:** 10.1101/406967

**Authors:** Rahmioglu Nilufer, Banasik Karina, Christofidou Paraskevi, Danning Rebecca, Galarneau Genevieve, Giri Ayush, MacGregor Stuart, Mortlock Sally, Sapkota Yadav, Schork J Andrew, Sobalska-Kwapis Marta, Stefansdottir Lilja, Turman Constance, Uimari Outi, Adachi Sosuke, Andrews Shan, Arnadottir Ragnheidur, Burgdorf S Kristoffer, Campbell Archie, Cheuk SK Cecilia, Clementi Caterina, Cook James, De Vivo Immaculata, DiVasta Amy, O Dorien, Edwards Todd, Fontanillas Pierre, Fung N Jenny, Geirsson T Reynir, Girling Jane, Harris R Holly, Holdsworth-Carson Sarah, Houshdaran Sahar, Hu-Seliger Tina, Jaqrvelin Marjo-Riitta, Kepka Ewa, Kulig Bartosz, Laufer R Marc, Law Matthew, Low Siew-Kee, Mangino Massimo, Marciniak Blazej, Matalliotaki Charoula, Matalliotakis Michail, Murray D Alison, Nezhat Camran, Nõukas Margit, Olsen Catherine, Padmanabhan Sandosh, Paranjpe Manish, Parfitt David-Emlyn, Peters Maire, Polak Grzegorz, Porteous J David, Romanowicz Hanna, Saare Merli, Shafrir Amy, Siewierska-Górska Anna, Skarp Sini, Slomka Marcin, Smith H Blair, Smolarz Beata, Szaflik Tomasz, Szyllo Krzysztof, Terry Kathyrn, Thorleifsson Gudmar, Tomassetti Carla, Vanhie Arne, Vincent Katy, Vitonis Allison, Werge Thomas, DBDS Genetic Consortium, S Andersen, K Banasik, S Brunak, KS Burgdorf, C Erikstrup, TF Hansen, H Hjalgrim, G Jemec, P Jennum, KR Nielsen, M Nyegaard, HM Paarup, OB Pedersen, M Petersen, E Sorensen, H Ullum, T Werge, D Gudbjartsson, K Stefansson, H Stefansson, U Þorsteinsdottir, iPSYCH-Broad Consortium, PB Mortensen, DM Hougaard, AD Borglum, the Celmatix Research Team, the 23andMe Research Team, Chasman I Daniel, D’Hooghe Thomas, Giudice C Linda, Goulielmos N George, Hapangama K Dharani, Hayward Caroline, Horne W Andrew, Kamatani Yoichiro, Kubo Michiaki, Martikainen Hannu, Rogers AW Peter, Saunders T Philippa, Sirota Marina, Spector Tim, Strapagiel Dominik, Tung Y Joyce, Whiteman David, Becker M Christian, Salumets Andres, Magi Reedik, Kraft Peter, Nyegaard Mette, Nyholt R Dale, Steinthorsdottir Valgerdur, Stefansson Kari, Velez-Edwards R Digna, Yurttas Beim Piraye, Missmer A Stacey, Montgomery W Grant, Morris P Andrew, Zondervan T Krina

## Abstract

Endometriosis is a common complex inflammatory condition characterised by the presence of endometrium-like tissue outside the uterus, mainly in the pelvic area. It is associated with chronic pelvic pain and infertility, and its pathogenesis remains poorly understood. The disease is typically classified according to the revised American Fertility Society (rAFS) 4-stage surgical assessment system, although stage does not correlate well with symptomatology or prognosis. Previously identified genetic variants mainly are associated with stage III/IV disease, highlighting the need for further phenotype-stratified analysis that requires larger datasets. We conducted a meta-analysis of 15 genome-wide association studies (GWAS) and a replication analysis, including 58,115 cases and 733,480 controls in total, and sub-phenotype analyses of stage I/II, stage III/IV and infertility-associated endometriosis cases. This revealed 27 genetic loci associated with endometriosis at the genome-wide p-value threshold (P<5×10^−8^), 13 of which are novel and an additional 8 novel genes identified from gene-based association analyses. Of the 27 loci, 21 (78%) had greater effect sizes in stage III/IV disease compared to stage I/II, 1 (4%) had greater effect size in stage I/II compared to stage III/IV and 17 (63%) had greater effect sizes when restricted to infertility-associated endometriosis cases compared to overall endometriosis. These results suggest that specific variants may confer risk for different sub-types of endometriosis through distinct pathways. Analyses of genetic variants underlying different pain symptoms reported in the UK Biobank showed that 7/9 had positive significant (p<1.28×10^3^) positive genetic correlations with endometriosis, suggesting a genetic basis for sensitivity to pain in general. Additional conditions with significant positive genetic correlations with endometriosis included uterine fibroids, excessive and irregular menstrual bleeding, osteoarthritis, diabetes as well as menstrual cycle length and age at menarche. These results provide a basis for fine-mapping of the causal variants at these 27 loci, and for functional follow-up to understand their contribution to endometriosis and its potential subtypes.

## Introduction

Endometriosis is an enigmatic chronic inflammatory condition occurring in 5-10% of women of reproductive age, associated with debilitating pelvic pain and infertility [1]. It is characterized by the presence of endometrial-like tissue outside the uterus, mainly on pelvic organs. Definitive diagnosis requires visualization through laparoscopy, and treatments are limited to surgical removal of disease, lysis of adhesions and hormonal treatments with often-intolerable side-effects. Endometriosis is a heterogeneous condition, typically classified according to rAFS criteria - with stage I/II disease featuring superficial peritoneal lesions and minimal adhesions, and stage III/IV more extensive disease including fibrosis, cystic ovarian endometriosis, deeply invasive nodules, and advanced scarring, fibrosis and adhesions [2]. Causes of endometriosis remain largely unknown, but the condition has an estimated heritability of ∼ 50% [3, 4] with ∼26% estimated to be due to common genetic variation [5]. In total eight genome-wide association studies (GWAS) in women of European and East Asian ancestry have been published to date [6], the largest of which comprised 17,045 cases and 191,596 controls and identified 19 distinct signals at 14 loci [7], leaving many loci yet to be uncovered. Although positional evidence had suggested potential involvement of sex-steroid hormone, WNT signalling, cell adhesion/migration, cell growth/carcinogenesis and inflammation-related pathways, the regulatory effects of most of these variants have yet to be determined in tissues relevant to endometriosis. Moreover, the causal variants of the identified signals and their functional involvement in mediating the underlying mechanisms remain yet to be uncovered.

Previous GWAS analyses have highlighted that the signals for most loci are driven by rAFS stage III/IV disease [2, 7]. However, it is unclear what phenotypic characteristics in this broad classification (e.g. cystic ovarian endometriosis, advanced scarring, fibrosis and adhesions) these associations pertain to, highlighting the need for deep phenotypically stratified analyses. More detailed sub-phenotypic analyses, based on infertility, pain symptomatology, and other surgical phenotypes could help decipher causal mechanisms contributing to various components of the complex underlying pathophysiology and ultimately more precise treatment targets.

Here, we have performed the largest GWAS and replication meta-analysis of endometriosis to date, including 58,115 cases and 733,480 controls. Moreover, we have conducted sub-phenotype analysis of surgically confirmed rAFS Stage I/II (3,160 cases), Stage III/IV disease (3,711 cases), as well as infertility-associated endometriosis (2,843 cases) vs. 482,225 controls (Table 1). We also investigated the shared genetic correlation of endometriosis with various previously reported co-morbid autoimmune, metabolic, reproductive and pain-related conditions [8-12].

**Table 1.** Characteristics of case and control sets from each participating centre.

| Study | Ancestry | Cases | Sub-phenotypes |  |  | Controls | Imputation Reference |
| --- | --- | --- | --- | --- | --- | --- | --- |
|  |  |  | Stage III/IV | Stage I/II | Infertile |  |  |
| BBJ | Japanese | 1,423 | - | - | - | 1,318 | 1000G P3v5 |
| iPSYCH | European | 397 | - | - | - | 9,385 | 1000G P3v5 |
| DECODE | European | 1,857 | 695 | 587 | - | 132,978 | deCODE |
| LEUVEN | European | 998 | 423 | 568 | - | 783 | 1000G P1 |
| NFBC | European | 168 | - | - | 55 | 9,194 | HRC |
| EGCUT | European | 1,716 | - | - | - | 35,176 | Estonian P |
| QIMRHCS | European | 2,262 | 905 | 1,354 | 848 | 2,923 | HRC |
| MELBOURNE | European | 320 | 78 | 242 | 31 | 887 | HRC |
| WGHS | European | 1,494 | - | - | - | 14,033 | HRC |
| NHS2 | European | 2,104 | - | - | 1,055 | 5,854 | 1000G P3v5 |
| Generation Scotland | European | 323 | - | - | - | 9,788 | HRC |
| UK Biobank | European | 6,611 | 1,075 | - | 417 | 251,704 | HRC |
| OXEGENE | European | 924 | 454 | 329 | 344 | 5,190 | HRC |
| Oxford-P1 | European | 165 | 81 | 80 | 93 | 78 | HRC |
| LODZ | European | 171 | - | - | - | 2934 | 1000G P3v5 |
| **Discovery Meta-analysis Total** |  | **20,933** | **3,711** | **3,160** | **2,843** | **482,225** |  |
| 23andMe Replication | European | 37,182 | - | - | - | 251,255 | 1000G P1v3 |
| **Combined Meta-analysis Total** |  | **58,115** | **3,711** | **3,160** | **2,843** | **733,480** |  |

## Results

Discovery GWAS meta-analysis included 15 studies composed of 20,933 cases and 482,225 controls of mainly European, and one study of East-Asian, ancestry. The replication analysis included 37,182 cases and 251,255 controls of European ancestry, bringing the total to 58,115 cases and 733,480 controls (Table 1). All individual datasets were imputed separately up to 1000 Genomes (1000G P3v5), Haplotype Reference Consortium (HRC r1.1 2016) or population-specific whole genome sequence data, for each ancestry (Table 1). A summary of each of the individual GWAS datasets is given in the Materials and Methods. Meta-analysis was conducted for 10,260,082 SNPs using the inverse weight variance method employed in METAL [13] for overall endometriosis and each sub-phenotype: Stage I/II, Stage III/IV and infertility-associated endometriosis. Post meta-analysis, we excluded variants that were not present in more than 50% of the effective sample size and retained 8,311,853 SNPs for analysis. For each locus associated at p<5×10^−5^, lead SNPs and their two proxies were followed up in replication analysis and meta-analysed across datasets (combined meta-analysis). Genome-wide significant loci were defined as containing SNPs associated at p<5×10^−8^ in the combined meta-analysis and p<7.14×10^−4^ (0.05/70 N lead SNPs) in the replication analysis. The QQ-plots and Manhattan plots for all analyses are provided in Supplementary Figures 1-5.

**Figure 1.**
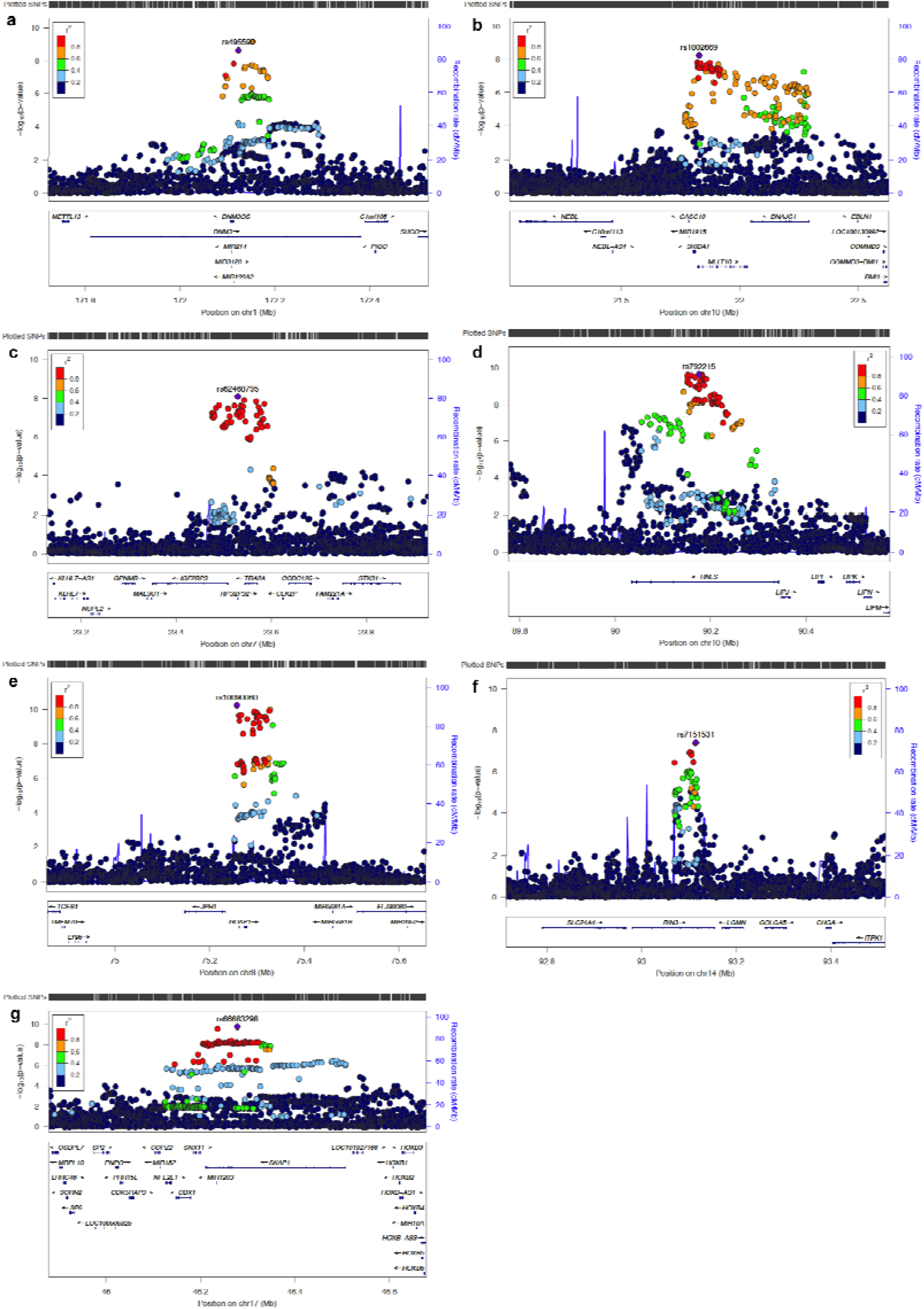
Regional association plots for the novel seven genome-wide significant endometriosis loci from discovery meta-analysis; (a) *DNM3 (1q24.3),* (b) *MIIT10 (10p12.31)* (c) *IGF2BP3 (7p15.3),* (d) *RNtS (10q23.31),* (e) *GDAP1 [8q21.II),* (f) *RIN3 (14q32.12),* (g) *SKAP1 (17q21.32).* The association results are shown on the y-axis as -logio(P-value) and on the x-axis is the genomic location (hg 19). The top associated SNP is coloured purple and the other SNPs are coloured according to the strength of LD with the top SNP by r in the European 1000 Genomes dataset.

### GWAS meta-analysis results: 13 novel loci for endometriosis

We identified 27 genome-wide significant loci for overall endometriosis, of which 13 were novel associations (Table 2, Supplementary Figures 1-2). Of these 13 novel loci, seven were identified in the discovery meta-analysis and the remaining six reached genome-wide significance when the replication data were folded into the combined meta-analysis. The novel signals of association from the discovery meta-analysis with overall endometriosis (Figure 1, Table 2) mapped to *DNM3* on 1q24.3 (rs495590 OR=1.07, 95% CI: 1.05-1.10, p=6.73×10^10^), near *IGF2BP3* on 7p15.3 (rs62468795 OR=1.10, 95% CI: 1.07-1.14, p=8.05×10^−9^), near *GDAP1* on 8q21.11 (rs10090060 OR=1.08, 95% CI: 1.06-1.11, p=5.72×10^−11^), in *MIIT10* on 10pl2.31 (rsl802669 OR= 1.07, 95% CI: 1.05-1.10, p=5.52×10^−9^), in *RNLS* on 10q23.31 (rs796945 OR=1.07, 95% CI: 1.05-1.10, p=1.78×10^−9^), in *RIN3* on 14q32.12 (rs7151531 OR=1.07, 95% CI: 1.04-1.10, p=3.80×10^−8^), and in *SKAP1* on 17q21.32 (rs66683298 OR=1.08, 95% CI: 1.06-1.11, p=l.73×10^10^). The replication analysis revealed six further novel loci (Table 2; Figure 2): rs1894692 in an intergenic regions between *SLC19A2* and *F5* on 1q24.2 (OR=1.18, 95% CI: 1.13-1.24 p=2.88×10^−13^) rs2510770 sits in *PDLIM5* on *4q22.3* (OR=1.05, 95% CI: 1.03-1.06 p=8.25×10^10^); rs13177597 near *ATP6AP1L* on 5q14.2 (OR=1.06, 95% CI: 1.04-1.08 p=1.30×10^−8^); rs17727841 in *IGF1* on *12q23.2* (OR=1.06, 95% CI: 1.04-1.08 p=5.33×10^−11^); rs4923850 near *BMF* on 15q15.1 (OR=1.05, 95% CI:1.04-1.06 p=3.07×10^−13^) and rs76731691 in *CEP112* on 17q24.1 (OR=1.08, 95% CI: 1.05-1.11 p=9.27×10^−9^).

**Table 2.** 27 genome-wide significant loci from the GWAS meta-analysis for endometriosis, stage III/IV, stage I/II and infertile endometriosis.

| Chr | Lead SNP | Position<br>(hg19) | RA (RAF) | Overall Endometriosis |  | Stage III/IV Endometriosis |  | Stage I/II Endometriosis |  | Infertile Endometriosis |  | Nearest Gene/<br>Cytoband |
| --- | --- | --- | --- | --- | --- | --- | --- | --- | --- | --- | --- | --- |
|  |  |  |  | OR (95% CI) | P value | OR (95% CI) | P value | OR (95% CI) | P value | OR (95% CI) | P value |  |
| Previously reported loci |  |  |  |  |  |  |  |  |  |  |  |  |
| 1 | rs12037376 | 22462111 | A (0.19) | 1.16 (1.13-1.19) | 2.85x10<sup>-22</sup> | 1.23 (1.15-1.32) | 1.33x10<sup>-9</sup> | 1.17 (1.08-1.26) | 7.43x10<sup>-5</sup> | 1.21 (1.11-1.31) | 4.24x10<sup>-6</sup> | WNT4/1p36.12 |
| 2 | rs11674184 | 11721535 | T (0.61) | 1.11 (1.09-1.14) | 1.47x10<sup>-18</sup> | 1.17 (1.11-1.23) | 1.13x10<sup>-8</sup> | 1.09 (1.02-1.16) | 6.82x10<sup>-3</sup> | 1.05 (0.99-1.12) | 0.11 | GREB1/2p25.1 |
| 2 | rs4141819 | 67864675 | C (0.30) | 1.05 (1.03-1.06) | 3.08x10<sup>-10*</sup> | 1.09 (1.03-1.15) | 1.53x10<sup>-3</sup> | 1.07 (1.00-1.14) | 0.03 | 1.06 (1.00-1.14) | 0.05 | ETAA1/2p14 |
| 2 | rs10167914 | 113529183 | C (0.32) | 1.08 (1.05-1.11) | 5.37x10<sup>-10</sup> | 1.13 (1.07-1.20) | 1.05x10<sup>-5</sup> | 1.06 (1.00-1.13) | 0.06 | 1.13 (1.05-1.19) | 5.61x10<sup>-4</sup> | IL1A/2q13 |
| 2 | rs1250247 | 216299629 | C (0.28) | 1.07 (1.05-1.10) | 3.88x10<sup>-8</sup> | 1.12 (1.06-1.18) | 6.82x10<sup>-5</sup> | 1.09 (1.02-1.16) | 9.67x10<sup>-3</sup> | 1.09 (1.02-1.16) | 0.01 | FN1/2q35 |
| 4 | rs10012589 | 56002689 | A (0.72) | 1.10 (1.07-1.13) | 5.69x10<sup>-13</sup> | 1.21 (1.14-1.28) | 1.86x10<sup>-10</sup> | 1.08 (1.01-1.15) | 0.02 | 1.11 (1.04-1.19) | 2.07x10<sup>-3</sup> | KDR/4q12 |
| 6 | rs6938760 | 19802104 | A (0.39) | 1.09 (1.06-1.11) | 2.89x10<sup>-12</sup> | 1.13 (1.08-1.19) | 1.88x10<sup>-6</sup> | 1.06 (1.00-1.13) | 0.05 | 1.10 (1.04-1.17) | 1.16x10<sup>-3</sup> | ID4/6p22.3 |
| 6 | rs7759516 | 151838245 | C (0.18) | 1.14 (1.11-1.17) | 1.35x10<sup>-18</sup> | 1.27 (1.19-1.35) | 6.58x10<sup>-13</sup> | 1.03 (0.96-1.12) | 0.41 | 1.08 (1.00-1.17) | 0.04 | CCDC170/6q25.1 |
| 6 | rs71575922 | 152554014 | G (0.16) | 1.15 (1.12-1.19) | 1.71x10<sup>-18</sup> | 1.30 (1.22-1.39) | 3.63x10<sup>-14</sup> | 1.10 (1.02-1.19) | 0.02 | 1.14 (1.05-1.23) | 2.36x10<sup>-3</sup> | SYNE1/6q25.1 |
| 7 | rs12700667 | 25901639 | A (0.72) | 1.08 (1.05-1.11) | 8.63x10<sup>-10</sup> | 1.20 (1.13-1.27) | 3.15x10<sup>-9</sup> | 1.06 (0.99-1.13) | 0.08 | 1.12 (1.05-1.20) | 5.75x10<sup>-4</sup> | 7p15.2/7p15.2 |
| 7 | rs55909142 | 46673774 | C (0.6) | 1.08 (1.05-1.10) | 8.76x10<sup>-11</sup> | 1.14 (1.08-1.20) | 1.61x10<sup>-6</sup> | 1.06 (1.00-1.13) | 0.05 | 1.11 (1.05-1.17) | 6.23x10<sup>-4</sup> | 7p12.3/7p12.3 |
| 9 | rs9987548 | 22173075 | A (0.4) | 1.09 (1.06-1.12) | 2.29x10<sup>-13</sup> | 1.19 (1.13-1.25) | 6.90x10<sup>-11</sup> | 1.05 (0.99-1.12) | 0.11 | 1.16 (1.09-1.24) | 7.14x10<sup>-7</sup> | CDKN2-BAS1/9p21.3 |
| 11 | rs74485684 | 30242287 | T (0.84) | 1.15 (1.12-1.19) | 1.32x10<sup>-18</sup> | 1.22 (1.14-1.31) | 3.54x10<sup>-8</sup> | 1.10 (1.02-1.20) | 0.01 | 1.17 (1.08-1.27) | 2.34x10<sup>-4</sup> | FSHB/11p14.1 |
| 12 | rs12320196 | 95712695 | C (0.3) | 1.08 (1.05-1.10) | 3.38x10<sup>-9</sup> | 1.14 (1.08-1.20) | 2.66x10<sup>-6</sup> | 1.09 (1.02-1.16) | 0.01 | 1.10 (1.03-1.17) | 4.15x10<sup>-3</sup> | VEZT/12q22 |
| Novel loci |  |  |  |  |  |  |  |  |  |  |  |  |
| 1 | rs1894692 | 169467654 | A (0.98) | 1.18 (1.13-1.24) | 2.88x10<sup>-13*</sup> | 1.63 (1.28-2.07) | 5.99x10<sup>-5</sup> | 1.12 (0.88-1.43) | 0.34 | 1.34 (1.06-1.71) | 0.02 | SLC19A2/1q24.2 |
| 1 | rs495590 | 172152202 | G (0.52) | 1.07 (1.05-1.10) | 6.73x10<sup>-10</sup> | 1.11 (1.06-1.17) | 3.22x10<sup>-5</sup> | 1.06 (1.00-1.12) | 0.05 | 1.07 (1.01-1.14) | 0.02 | DNM3/1q24.3 |
| 4 | rs2510770 | 95479372 | A (0.33) | 1.05 (1.03-1.06) | 8.25x10<sup>-10*</sup> | 1.06 (1.01-1.12) | 0.03 | 1.06 (0.99-1.13) | 0.08 | 1.08 (1.01-1.15) | 0.02 | PDLIM5/4q22.3 |
| 5 | rs13177597 | 82052282 | G (0.13) | 1.06 (1.04-1.08) | 1.30x10<sup>-8*</sup> | 1.15 (1.06-1.24) | 6.81x10<sup>-4</sup> | 1.07 (0.97-1.17) | 0.16 | 1.11 (1.01-1.22) | 0.04 | ATP6AP1L/5q14.2 |
| 7 | rs62468795 | 23530051 | G (0.85) | 1.10 (1.07-1.14) | 8.05x10<sup>-09</sup> | 1.14 (1.06-1.22) | 5.2x10<sup>-4</sup> | 1.15 (1.06-1.26) | 1.24x10<sup>-3</sup> | 1.08 (0.99-1.18) | 0.08 | IGF2BP3/7p15.3 |
| 8 | rs10090060 | 75257608 | A (0.58) | 1.08 (1.06-1.11) | 5.72x10<sup>-11</sup> | 1.08 (1.02-1.14) | 3.85x10<sup>-3</sup> | 1.09 (1.02-1.15) | 6.80x10<sup>-3</sup> | 1.11 (1.05-1.18) | 4.83x10<sup>-4</sup> | GDAP1/8q21.11 |
| 10 | rs1802669 | 21827796 | A (0.34) | 1.07 (1.05-1.10) | 5.52x10<sup>-9</sup> | 1.09 (1.03-1.15) | 2.14x10<sup>-3</sup> | 1.09 (1.03-1.16) | 6.08x10<sup>-3</sup> | 1.09 (1.02-1.16) | 6.41x10<sup>-3</sup> | MLLT10/10p12.31 |
| 10 | rs796945 | 90150837 | C (0.37) | 1.07 (1.05-1.10) | 1.78x10<sup>-9</sup> | 1.08 (1.03-1.14) | 2.83x10<sup>-3</sup> | 1.05 (0.99-1.12) | 0.10 | 1.11 (1.05-1.18) | 5.80x10<sup>-4</sup> | RNLS/10q23.31 |
| 12 | rs17727841 | 102809630 | G (0.81) | 1.06 (1.04-1.08) | 5.33x10<sup>-11*</sup> | 1.17 (1.09-1.25) | 4.95x10<sup>-6</sup> | 1.06 (0.98-1.14) | 0.16 | 1.03 (0.95-1.11) | 0.46 | IGF1/12q23.2 |
| 14 | rs7151531 | 93113547 | C (0.29) | 1.07 (1.04-1.10) | 3.80x10<sup>-8</sup> | 1.10 (1.04-1.16) | 8.37x10<sup>-4</sup> | 1.01 (0.95-1.08) | 0.76 | 1.11 (1.04-1.18) | 1.82x10<sup>-3</sup> | RIN3/14q32.12 |
| 15 | rs4923850 | 40352278 | A (0.47) | 1.05 (1.04-1.06) | 3.07x10<sup>-13*</sup> | 1.10 (1.04-1.15) | 4.42x10<sup>-4</sup> | 1.03 (0.97-1.09) | 0.40 | 1.07 (1.01-1.13) | 0.03 | BMF/15q15.1 |
| 17 | rs66683298 | 46277748 | C (0.61) | 1.08 (1.06-1.11) | 1.73x10<sup>-10</sup> | 1.10 (1.04-1.16) | 3.71x10<sup>-4</sup> | 1.10 (1.04-1.17) | 1.93x10<sup>-3</sup> | 1.06 (1.00-1.13) | 0.04 | SKAP1/17q21.32 |
| 17 | rs76731691 | 63960269 | G (0.94) | 1.08 (1.05-1.11) | 9.27x10<sup>-9*</sup> | 1.20 (1.08-1.34) | 9.77x10<sup>-4</sup> | 1.25 (1.10-1.42) | 7.29x10<sup>-4</sup> | 1.14 (1.01-1.29) | 0.04 | CEP112/17q24.1 |

\* The overall endometriosis association results are from the combined meta-analysis including replication evidence that unrevealed genome-wide significant results. Discovery overall endometriosis GWAS meta-analysis results for these variants were: rs4141819 p=9.5x10<sup>-8</sup>, OR=1.06, 95% CI=1.04-1.09; rs1894692 p=4.18x10<sup>-6</sup>, OR=1.22, 95% CI=1.12-1.33; rs2510770 p=1.03x10<sup>-6</sup>, OR=1.06, 95% CI=1.04-1.09; rs13177597 p=9.9x10<sup>-6</sup>, OR=1.08, 95% CI=1.05-1.12; rs17727841 p=6.09x10<sup>-8</sup>, OR=1.08, 95% CI=1.05-1.12; rs4923850 p=1.06x10<sup>-7</sup>, OR=1.06, 95% CI=1.04-1.09; rs76731691 p=1.06x10<sup>-6</sup>, OR=1.13, 95% CI=1.07-1.18.

**Figure 2.**
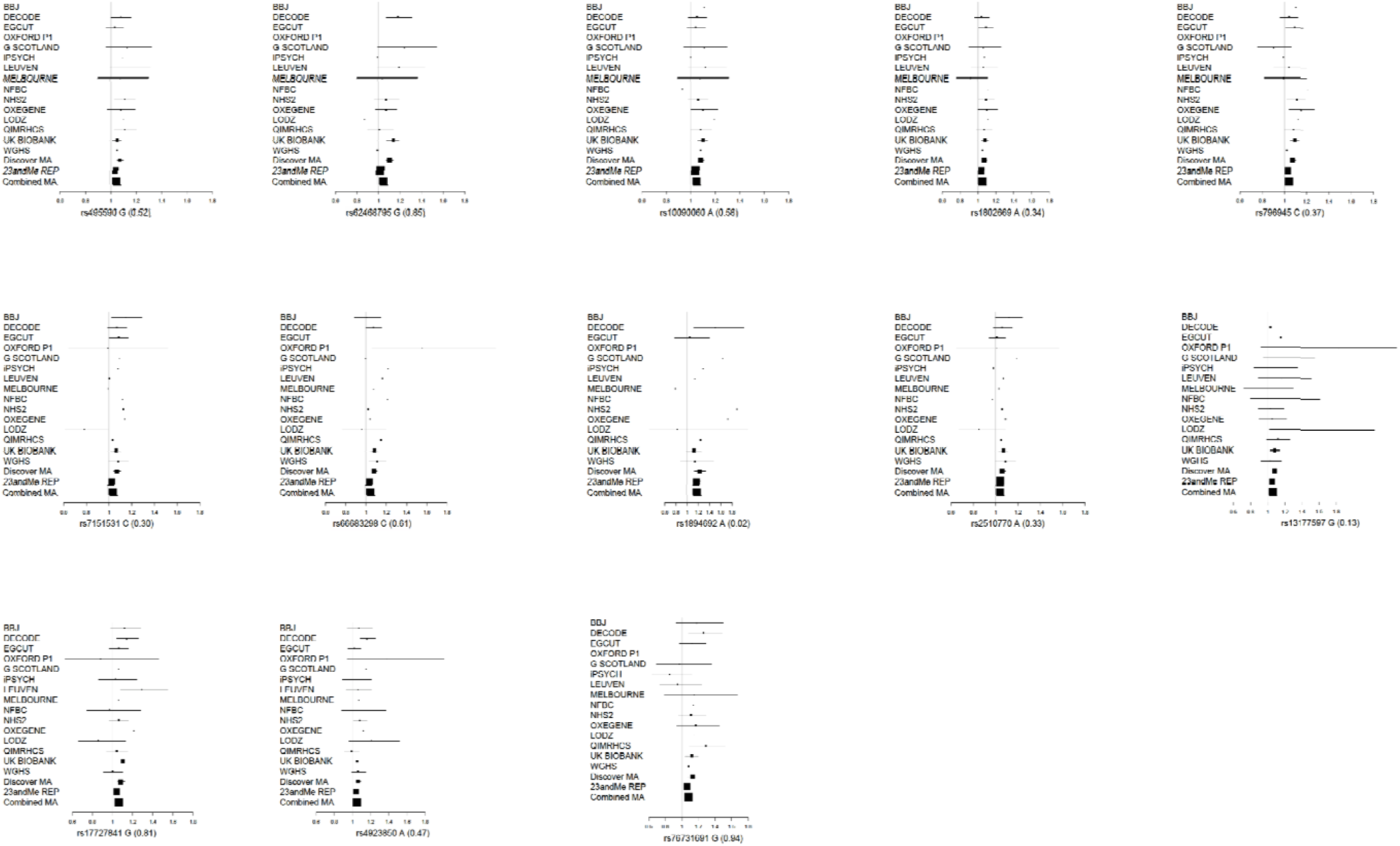
Forest plots for the lead SNPs of 13 novel genome-wide significant endometriosis loci (combined meta-analysis). The rsid and the effective allele along with effective allele frequency are given under each plot. The x-axis displays the association odds-ratio scale, and association results from each study are presented with study acronym. The discovery meta-analysis results (Discovery MA), 23andMe replication results (23andME REP) and combined meta-analysis results (Combined MA) are also illustrated.

In the sub-phenotype GWAS there were eight genome-wide significant signals associated with stage III/IV disease and one borderline genome-wide significant signal when cases were restricted to infertility-associated endometriosis, rs7041895 near *CDKNB-AS1* on 9p21.3 (OR=1.18, 95% CI: 1.11-1.25, p=6.89×10^−8^), all of which were also genome-wide significant for overall endometriosis (Table 2).

Of all 27 loci, 21 had larger effect sizes for stage III/IV compared with stage I/II disease *(SLC19A1, DNM3, ATP6AP1L, RNLS, IGF1, RIN3, BMF, WNT4/CDC42, GREB1, ETAA1, IL1A, FN1, KDR, ID4, CCDC170, SYNE1, 7pl5.2, 7pl2.3, CDKN2BAS1, FSHB, VEZT)* and 8/21 were also genome-wide significant in stage III/IV disease; five loci had similar effect sizes with both surgical stages (*PDLIM5, IGF2BP3, GDAP1, MIIT10, SKAP1);* and one locus had larger effect sizes for stage I/II vs. stage III/IV (*CEP112).* Infertility-associated endometriosis cases showed larger effect sizes compared with overall endometriosis for 17 loci (.*SLC19A2, PDLIM5, ATP6AP1L, GDAP1, MIIT10, RNLS, RIN3, BMF, CEP112, WNT4/CDC42, IL1A, FN1, 7pl5.2, 7pl2.3, CDKN2-BAS1, FSHB, VEZT);* similar for 5 loci *(DNM3, ETAA1, KDR, ID4, SYNE1)* and decreased for 5 loci (*IGF2BP3, IGF1, SKAP1, GREB1, CCDC170)* (Table 2, Supplementary Figures 6i-vii and 7i-xiii).

Among the 13 novel loci, the lead SNP (rs495590) in *DNM3* showed an association with stage III/IV disease (stage III/IV OR=l.11, 95% Cl: 1.06-1.17 p=3.22×10^−5^) but only marginally with stage I/II (OR=1.06, 95% CI: 1.00-1.12, p=0.05) or infertility-associated endometriosis (OR=1.07, 95% CI: 1.01-1.14, p=0.02) (Table 2, Supplementary Figure 6i). The lead SNP maps to an intron within the *DNM3* (Dynamin 3) that encodes for a GTP-binding protein that is associated with microtubules and involved in vesicular transport.

Both rs62468795 on 7p15.3 and the intronic rs66683298 in *SKAP1* (Src Kinase Associated Phosphoprotein 1) on 17q21.32 showed nearly identical effect estimates with stage I/II and stage III/IV (OR_stageI/II_=l-15, 95% Cl: 1.06-1.26; OR_stageIII/IV_=l-14, 95% Cl: 1.06-1.22 and OR_stageI/II_=1.10 95% CI:1.04-1.17; OR_stageIII/IV_=1.10 95% 0:1.04-1.16, respectively), but they were not associated with infertile endometriosis (OR=1.08, 95% CI: 0.99-1.18, p=0.08; OR=1.06, 95% CI: 1.00-1.13, p=0.04) relative to all controls regardless of infertility status (Table 2, Supplementary Figures 6iii and 6vii). rs62468795 is located 20.1kb downstream of *IGF2BP3* (Insulin Like Growth Factor 2 MRNA Binding Protein 3), 14.35kb upstream of *TRA2A* (Transformer 2 Alpha Homolog), and 0.98kb upstream of *RPS2P32,* a non-coding RNA. *IGF2BP3* is an RNA binding factor that is involved in regulation of translation of insulin like growth factor II. It also promotes cell adhesion and protrusion of plasma membrane associated with degradation of extracellular matrix in cancer invasiveness and metastasis.

*TRA2A* is involved in regulation of pre-mRNA splicing. rs66683298 sit in an intron in *SKAP1* and there is strong LD covering upstream of *SKAP1* which also harbours *SNX11, CBX1* and *NFE2L1* (Figure 1g). *SKAP1* has immuno-regulatory functions including, regulation of T-cell receptor signalling by enhancing the MAP kinase pathway, and optimisation of conjugation between T-cells and antigen-presenting cells.

rs10090060 on 8q21.11 showed a stronger association with infertile endometriosis 0R_infertile_^=^ 1.11, 95% CI: 1.05-1.18, p=4.83×10^−4^ vs. OR_Overall_=1.08 95%CI: 1.05-1.15, p=5.72×10^−11^) (Table 2, Supplementary Figure 6v). It sits 5.01kb upstream of *GDAP1* (Ganglioside Induced Differentiation Associated Protein 1) with strong LD coverage of the *GDAP1* gene. *GDAP1* plays a role in neuronal development and is likely to be related to pain processing.

rs1802669 is an intronic variant located in *MIIT10* (Histone Lysine Methyltransferase DOT1L Cofactor) on 10p12.31. It did not show differential association with any of the sub-types of endometriosis (Table 2, Supplementary Figure 6ii). *MIIT10* is a transcription factor involved in various chromosomal rearrangements resulting in different types of leukemia.

rs796945 and rs7151531 are intronic SNPs located in *RNLS* (Renalase, FAD Dependent Amine Oxidase) on 10q23.31 and *RIN3* (Ras And Rab Interactor 3) on 14q32.12, respectively. They both have larger effect sizes for stage III/IV disease compared with stage I/II disease (OR_stageIII/IV_=1.08, 95% CI: 1.03-1.14, p=2.83×10^−3^; OR_stageIII/IV_^=^ 1 -10, 95% Cl: 1.04-1.16, p=8.37×10^−4^ vs. OR_stageI/II_=1.05, 95% CI: 0.99-1.12, p=0.10; OR_stageI/II_=l-01, 95% Cl: 0.95-1.08, p=0.76) and infertility-associated endometriosis sub-group compared with overall endometriosis (OR_infertile_=l-H, 95% Cl: 1.05-1.18, p=5.8×10^−4^; OR_infertile_=l.II, 95% Cl: 1.04-1.18, p=1.82×10^−3^ vs. OR_overall_=1.07, 95% CI: 1.05-1.10, p=1.78×10^−9^ and OR_Overa11_=1.07, 95% CI: 1.04-1.10, p=3.80×10^−8^) (Table 2, Supplementary Figure 6iv and vi). *RNLS* encodes for an enzyme hormone secreted by kidney that catalyses the conversion of 1,2-dihydro-beta-NAD(P) to 1,6-dihydro-beta-NAD(P) to form beta-NAD(P)+. It circulates in blood modulating cardiac function and systemic blood pressure. *RIN3* encodes for a RAS effector protein that aids in functioning of *RAB5B* and *RAB31* oncogenes.

The six novel loci identified in the replication analysis of overall endometriosis were

compared to those identified by the discovery meta-analysis of endometriosis subphenotype. Four of these were entirely limited to the stage III/IV disease including: (1) rs1894692 in an intergenic regions between *SLC19A2* and *F5* on 1q24.2 which are involved in cellular transport and regulation of hemostasis respectively; (2) rs13177597 in an intergenic region with nearest gene named *ATP6AP1L* involved in cellular transport on 5q14.2; (3) rs17727841 in *IGF1* is involved in growth and development *on 12q23.2; (4)* rs4923850 sits near *BMF,* which is involved in regulation of apoptosis on 15q15.1 (Table 2).

rs2510770 did not show any differential association between any of the subphenotypes (Table 2) and it resides in *PDLIM5* known to be involved in heart development on *4q22.3.* rs76731691 was nominally associated with both surgical stage groups of endometriosis but with a larger effect in stage I/II disease and no association with infertile endometriosis (Table 2). This variant sits in *CEP112, which* encodes for a cell division control protein on 17q24.1.

Of all previously reported endometriosis GWAS loci, two were not replicated. They originally had been reported from a relatively small European ancestry GWAS of 171 cases and 2934 controls [14]: rs10129516 near *RHOJ* on 14q23.2 (OR=1.01, 95 CI: 0.98-1.05, p=0.59), and rs644045 near *C2* on 6p21.33 (OR=1.01, 95% CI: 0.98-1.03, p=0.56).

### Genes associated with endometriosis

A genome-wide gene-based analysis in MAGMA as implemented in FUMA [15] identified 34 genes surviving the genome-wide significance threshold of 2.67×10^−6^ (Supplementary Figure 8, Supplementary Figure 9, results with p<0.05 in Supplementary Table 2). Of these 34, 25 overlap with or are located near genome-wide significant loci including 1p36.12 (*WNT4, CDC42),* 1q24.3 (*DNM3*), 2p25.1 (*GREB1*), 2q13 *(IL1A),* 2q35 (F/VI), 6q25.1 *(CCDC170, ZBTB2, RMND1, C6orf211, ESR1, SYNE1),* 7p15.2 (*RP1-170019*), 8q21.11 (*GDAP1*), 10p12.31 (*MIIT10, DNAJC1),* 12q23.2 (*IGF1, NUP37, PARBP),* 10q23.31 *(RNLS),* IIp14.1 (*ARL14EP*), 12q22 *[VEZT, FGD6),* 14q32.12 *(RIN3),* 15q15.1 *{BMF),* 17q21.32 (*SKAP1*). The other nine genes were in eight novel genomic regions including 3p25.3 *{ATG7),* 6p21.31 (*HMGA1*), 6q13 (*CD109*), 6q22.33 (*RSP03*), 7q21.12 *(ADAM22),* 8q23.3 *(TRPS1),* 10p12.31 *{SKIDAI),* 12q13.13 (*HOXC6, RP11-834C11.12).* Of the nine novel genes, *ATG7* is involved in autophagy and vacuole transport and has been previously associated with increased HDL cholesterol levels *[16]; HMGA1* is involved in regulation of gene transcription, metastatic progression of cancer cells and is associated with Type 2 diabetes and fat deposition around the waist (waist circumference) [17]; *HOXC6* is also associated with fat deposition around the waist; *CD109* encodes for a glycoprotein found on the surfaces of activated T-cells and it has been associated with squamous cell carcinoma of the vulva. *RSP03* is another fat distribution associated gene that is involved in regulation of the *WNT*signalling pathway. Moreover, both *RSP03* and *HMGA1* are also associated with uterine fibroids [18]. *ADAM22* is involved in regulation of cell adhesion and proliferation. *TRPS1* encodes for a transcription factor that represses GATA-regulated genes and it was previously associated with monocyte percentage of white blood cells.

### Genetic enrichment between endometriosis and other traits and conditions

Pheno Scanner V1.1 [19] was utilised to investigate whether the 27 genome-wide significant lead SNPs from our endometriosis analysis were also associated with phenotypes previously established to be associated with endometriosis (r^2^>0.6 between endometriosis and other phenotype lead SNPs and p<0.001,Supplementary Table 3). This has revealed three genome-wide significant associations including: (1) rs12700667 on 7p15.2 with waist to hip ratio adjusted for body mass index (WHRadjBMI), (2) rs1894692 on 1q24.2 with venous thrombosis, (3) rs495590 on 1q24.3 with height. In addition, some interesting nominal genetic associations were identified: (1) Auto-immune conditions: rs1894692 on 1q24.2 with inflammatory bowel disease and rs7151531 on 14q32.12 with rheumatoid arthritis;(2) rs4141819 on 2p14 and rs17727841 on 12q23.2 with age of menopause, (3) rs796945 on 10q23.31 with age of menarche, (4) rs4141819 on 2p14, rs1250247 on 2q13 and rs17727841 on 12q23.2 with WHR, (5) rs1802669 on 10p12.31 and rs796945 on 10q23.31 with years of educational attainment, (6) rs1250247 on 2q13 and rs2510770 on 4q22.3 with coronary artery disease (Supplementary Table 3). Investigation in the NHGRI GWAS catalogue [20] with the nearest genes of the genome-wide significant endometriosis SN Ps revealed that eight of the genome-wide significant loci have been previously associated with breast cancer *(RIN3, MIIT10, CCDC170, ESR1, ATP6AP1L)* and/or ovarian cancer *(WNT4, MIIT10, SKAP1)* and pancreatic cancer (*IGF2BP3*), prostate cancer (*PDLIM5*) and colorectal *(CDKN2B-AS1, CDC42)* or endometrial cancer (*CDKN2B-AS1, BMF).* Multiple endometriosis loci were also associated with bone mineral density (*WNT4, ESR1, CCDC170, CEP112),* anthropometric measures including BMI, WHR, height and birth weight (*WNT4, DNM3, ESR1, NFE2L3, IGF2BP3, GDAP1, FSHB, IGF),* reproductive traits/conditions including sex hormone levels, polycystic ovary syndrome, menarche age at onset, menopause age at onset, length of menstrual cycle, gestational age at birth (’*WNT4/CDC42, E5R1, FSHB).* Also multiple loci were associated with coronary artery diseases (*FN1, CDKN2B-AS1, PDLIM5). GDAP1* has been associated with the intensity of dysmenorrhoea, whilst *RNLS* has been associated with type 1 diabetes and depressive/manic episodes in bipolar disorder (Supplementary Table 4).

To investigate genome-wide genetic overlap between endometriosis and various co-morbid autoimmune, metabolic, reproductive and pain-related conditions [8-12] and traits related to the biology of endometriosis, such as length of menstrual cycle, age of menarche, age at menopause, reproductive behavioural traits [21] and anthropometric measurements [22], we performed LD-score regression analysis as implemented in LD Hub using the featured publicly available GWAS datasets [23]. Out of the 39 traits/conditions included in the analysis, conditions including uterine fibroids, excessive and irregular menstrual bleeding, osteoarthritis, diabetes and dorsalgia showed significant (p<0.05/39=1.28×10^3^) positive genetic correlation with endometriosis (Figure 3). Irregular and excessive bleeding was most strongly correlated with stage III/IV disease. Osteoarthritis, a chronic inflammatory pain condition, showed the strongest genetic correlation with infertile endometriosis followed by stage III/IV disease. Uterine fibroids (leiomyoma of uterine smooth muscle) showed a strong positive genetic correlation with endometriosis regardless of disease sub-type. None of the autoimmune conditions show significant genetic correlation with endometriosis, despite previous reports of phenotypic correlation in epidemiological studies [24, 25] (Supplementary Figure 10).

**Figure 3.**
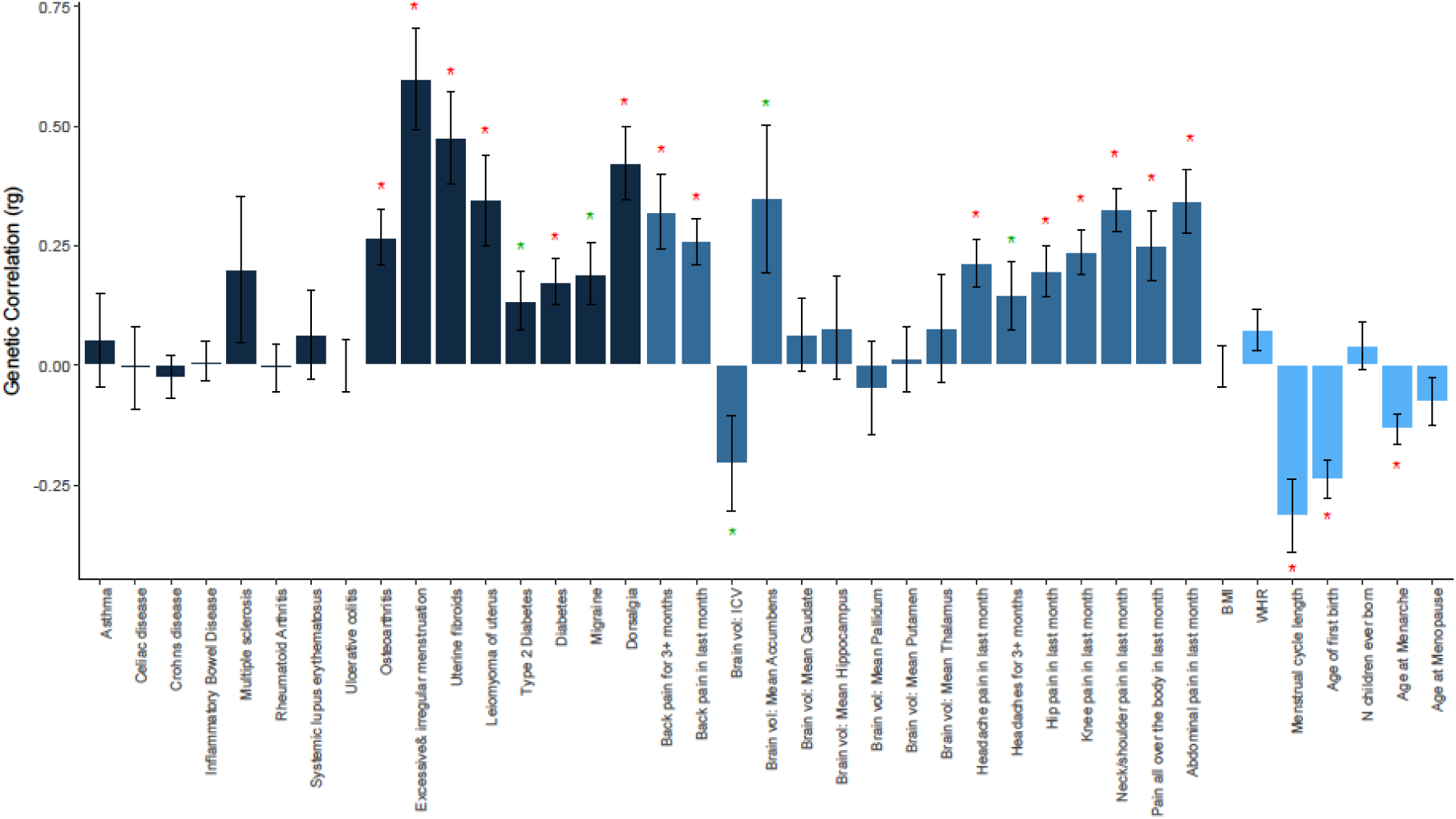
LD Score regression results for genetic enrichment between endometriosis and reproductive traits, anthropometric traits, pain-related symptomatology, and reproductive, autoimmune, metabolic and inflammatory conditions. The y-axis shows the genetic correlation (rg) between each condition and endometriosis with standard error bars. The x-axis shows the results per trait/condition. A red star denotes significant genetic enrichment after multiple-testing correction (p<l.28×10^−3^) and a green star nominal association (p<0.05).

The relationship between pain and endometriosis is complex. There is little correlation between pain symptoms and burden of disease and some women with endometriosis experience no pain at all. It has therefore been proposed that, particularly in the case of mild disease, those women vulnerable to pain may be more likely to experience pain [26, 27]. Additionally, the experience of dysmenorrhea and other painful symptoms associated with endometriosis has been shown to sensitise the nervous system [26, 28-30] potentially increasing the risk of other pain conditions. We therefore investigated the genetic correlation between endometriosis and nine different pain symptoms available from UK Biobank, of which seven showed significant positive genetic correlation with endometriosis (Figure 3). This illustrates that there is a shared genetic basis for the increased risk of other chronic pain conditions in patients with endometriosis pain. Moreover, Intracranial brain volume (ICV) of regions known to be associated with the perception of pain showed the strongest negative genetic correlation with infertility-associated endometriosis group and caudate brain volume showed strong positive genetic correlation with stage III/IV and infertile endometriosis sub-groups (Supplementary Figure 11).

Finally, menstrual cycle length, age at menarche onset and age of first birth showed significant negative genetic correlation with endometriosis (Figure 3). Earlier age of menarche is a risk factor for endometriosis [31], suggested to act through an increased exposure to oestrogen levels earlier in life. These analyses suggest this effect can be attributed to shared underlying genetic components or that age of menarche is causally related to endometriosis. Interestingly, number of children ever born was only negatively genetically correlated with stage III/IV disease (r=-0.23, p=5.6×10^−3^), but not with stage I/II disease (r=0.05, p=0.65) nor overall endometriosis (r=0.04, p=0.43) (Supplementary Figure 12). Age of first birth showed nominal negative genetic correlation with stage I/II disease (r=-0.23, p=0.02) and infertile endometriosis (r=-0.20, p=0.03) but not with stage III/IV disease (r=-0.04,p=0.56). Age at first birth and number of children ever born are both complex reproductive behaviour traits with heritability estimates up to 50%, 15% and 10% of variance due to common genetic variants respectively [21].

### Variance explained

The sibling relative risk *(λ*_*s*_) attributable to lead SNPs from known loci (N=14) rose from *λ*_*s*_*=* 1.026 to *λ*_*s*_*=* 1.037 after inclusion of the 13 novel loci. Assuming a population prevalence of 8% for endometriosis, the 14 known loci explained 1.52% of variance and including the novel 13 loci, all 27 loci together explained 2.15% of variance in disease susceptibility [32]. For stage III/IV disease, the sibling relative risk attributable to previously known loci rose from *λ*_*s*_ = 1.078 to *λ*_*s*_*=* 1.105 after inclusion of the novel loci. Assuming a population prevalence of 2.5% for stage III/IV endometriosis, the variance explained increased from 2.85% to 3.83%.

In conclusion, the findings show that the genetic underlying mechanisms of endometriosis implicate metabolic, reproductive, inflammatory and pain-processing/sensitivity pathways. Moreover, the variants that show stronger associations with infertile endometriosis or stage III/IV endometriosis further strengthens the fact that specific variants may confer risk for different sub-types of endometriosis through distinct pathways. These findings set the scene for fine-mapping the causal variants for each of the 27 loci and functional follow-up of identified variants to understand the underlying causal mechanisms for endometriosis as a single complex condition or a condition with multiple subtypes.

## Materials and Methods

### Subjects, Genotyping, Quality Control and Imputation

The meta-analysis included 15 datasets, which in total contribute 20,933 cases, and 482,225 controls of mainly European ancestry and one dataset from Japanese ancestry (1,423 cases: 1,318 controls). The datasets were genotyped in various genotyping arrays and each dataset was subject to sample and variant quality controls.

### BBJ

All Japanese GWAS case and control samples were obtained from BioBank Japan at the Institute of Medical Science at the University of Tokyo. A total of 1,423 cases were diagnosed with endometriosis by the presence of multiple clinical symptoms, physical examinations and/or laparoscopy or imaging tests. We used 1,318 female control samples from healthy volunteers from the Osaka-Midosuji Rotary Club (Osaka, Japan) and women in BioBank Japan who were registered to have no history of endometriosis. All participants provided written informed consent to this study. The study was approved by the ethical committees at the Institute of Medical Science at the University of Tokyo and the Center for Genomic Medicine at the RIKEN Yokohama Institute. The BBJ cases and controls were genotyped using the Illumina HumanHap550v3 Genotyping BeadChip. Quality control filtering required sample call rate of ≥0.98, IBS analysis was used to exclude samples with close relatedness and principal-component analysis was used to exclude non-Asian samples. We also performed SNP quality control (call rate of ≥0.99 in both cases and controls and HWE *P* of ≥1 × 10^−6^ in controls). In total, 460,945 SNPs on all chromosomes passed the quality control filters and followed up with imputation up to 1000G P3v5.

### DECODE

Endometriosis cases and controls from deCODE were from the deCODE Genetics, Iceland. Genome-wide imputation of this cohort was based on whole genome sequencing of 15,220 Icelanders using Illumina technology. Sequenced variants were then imputed into 151,677 Icelanders who had been genotyped using Illumina genotyping arrays. Using genealogic information, the sequence variants were also imputed into 282,894 un-typed relatives of chip-typed individuals to further increase the sample size for association analysis and increase the power to detect associations. Of these, this cohort included 1,857 surgically diagnosed endometriosis cases and 129,016 controls, with 695 cases exhibiting rAFS stage III/IV disease and 587 with rAFS stage I/II disease.

### EGCUT

The Estonian Biobank cohort is a volunteer-based sample of the Estonian resident adult population (aged ≥18 years). The current number of participants is close to 52000 and it represents a large proportion, 5%, of the Estonian adult population. At baseline, the general practioneers performed a standardized health examination of the participants, who also donated blood samples for DNA, white blood cells and plasma tests and filled out a 16-module questionnaire on health-related topics such as lifestyle, diet and clinical diagnoses described in WHO ICD-10 [33]. Endometriosis cases have been selected based on clinical diagnoses from the biobank questionnaire and from the latter linking of data to electronic health records. Additional set of endometriosis cases are recruited by prof. Andres Salumets from the patients from Tartu University Clinics. For these samples additional phenotypic information is available including endometriosis stage information.

For the genotype QC, following filters were applied call-rate>95%, HWE p-value>l × 10^−6^and MAF>1% prior the imputation. Samples were filtered by: MAF>1%, gender mismatches, heterozygosity<mean+-3SD. We have used Estonian population specific imputation reference panel [34] for imputation: Eagle v2.4 software for prephasing and Beagle 4.1 for the imputation. Association testing was done using EPACTS v.3.3 with Firth test using 10 first PC to adjust for population stratification.

### Generation Scotland

Generation Scotland: Scottish Family Health Study (GS:SFHS) is a family-based population cohort with DNA, biological samples, socio-demographic, psychological and clinical data from approximately 24,000 adult volunteers across Scotland. Although data collection was crosssectional, GS:SFHS became a prospective cohort due to of the ability to link to routine Electronic Health Record (EHR) data. Endometriosis cases were identified using the ICD9 (617) and ICD10 (N80) codes which resulted in 323 cases and 9788 female population controls. Genotyping was performed using HumanOmniExpressExome8v1_A and HumanOmniExpressExome8v1-2_A platforms. Standard quality control was conducted including removal variants with call rate>=97%, HWE *P* < 1 × 10^−6^ and MAF<0.01%. Post-QC 602451 markers were utilised in pre-phasing and imputation. Pre-phasing was conducted in using SHAPEIT v2 and data was imputed up to HRC rl.l2016. Association testing was conducted using Regscan [35] followed by correction of the association beta and standard error values in SNPTEST.

### iPSYCH2016

A total of 397 endometriosis cases and 9385 controls were drawn from the iPSYCH population-based dataset of more than 80,000 samples derived from the Danish Neonatal Screening Biobank. In short the Danish Neonatal Screening Biobank comprise dried bloodspots (Guthrie cards) from all individuals born in Denmark since 1981, where DNA extracted from the bloodspots can be successfully amplified and employed in GWAS [36]. All samples can be linked to the Danish register system, including the Danish National Patient Register (DNPR) that contains information about all hospital admissions and discharge, diagnoses and surgical procedures since 1977 [37]. The 80,000 iPSYCH samples were drawn primarily to study mental disorders, including approximately 30,000 population-based controls. The endometriosis cases were identified from the entire iPSYCH sample using ICD-8: 625.30-39 and ICD-10: N80.0-N80.9 (the 9th revision of ICD was not used in Denmark) from the DNPR (DNPR freeze ultimo 2016). The control individuals were selected to be age-matched individuals from the iPSYCH controls. All endometriosis cases in the iPSYCH sample were younger than 35 years. This study has been approved by the Danish research ethical committee system.

### Leuven

Cases (n = 1,077) and controls (n = 900) included in this study were recruited at the Leuven University Hospital, Belgium during 1993 - 2012, who had undergone laparoscopy for subfertility with or without pain. Presence of endometriosis in cases was laparoscopically and histologically confirmed based on electronic medical file records. The disease severity in women with endometriosis was prospectively graded according to the rAFS classification system. Endometriosis cases had either minimal (stage I; n = 380), mild (stage II, n = 229), moderate (stage III, n = 174), severe (stage IV; n = 284) or unknown (n = 10) disease. Absence of endometriosis in controls was confirmed laparoscopically. Both cases and controls were European in origin. All the study participants provided written informed consent and the study was approved by the Commission of Medical Ethics of the Leuven University Hospital, Belgium and QIMR Berghofer Human Ethics Research Committee, Australia. Whole genome genotyping of the DNA samples was performedusing the lllumina HumanCoreExome 12vl.l array at the Molecular Epidemiology Laboratory, QIMR Berghofer Medical Research Institute, Brisbane, Australia, following manufacturer’s standard protocol.

Genotype data were called using a custom cluster file generated using ∼2,000 good quality (<1% missing rate) samples and the GenCall algorithm within Illumina Genome Studio. Data were then further processed by zCall [38], a rare variant caller, to attempt to re-call missing genotypes. following guidelines by the manufacturer and protocols developed for Exome chip data, quality control (QC) measures were applied to the Leuven genome-wide association data. Briefly, samples with > 1% missing rates, outlying heterozygosity, non-European ancestries based on 1000 Genomes European populations, cryptic relatedness (pi-hat > 0.2) and gender discordances were excluded. Similarly, markers with poor separation of three genotype clusters, excess heterozygosity, outlying mean theta and intensity values for heterozygote genotypes, > 1% missing rates, Hardy-Weinberg Equilibrium (HWE) *P* < 10^−6^ in controls, and minor allele frequency (MAF) < 0.05% in cases and in controls were dropped. Following the QC steps, a total of 998 endometriosis cases (423 stage III/IV cases) and 783 disease-free controls remained in the Leuven genome-wide association data, which were imputed up to the 1000 Genomes Project reference panel (March 2012 release), using SHAPEIT [39] and minimac programs [40, 41] and following the two-step approach outlined in the online Minimac: 1000 Genomes Imputation Cookbook (http://genome.sph.umich.edu/wiki/Minimac:1000 Genomes Imputation Cookbook). Quality of the imputed genotypes was assessed by *R*^*^2^*^ metric, which estimates the squared correlation between true and imputed genotypes. Poorly imputed SNPs indicated by *R*^*^2^*^*<* 0.3 were excluded from the downstream analyses. Association analyses using the imputed genotype dosage scores were conducted using Plink for overall and stage III/IV endometriosis cases separately.

### LODZ-Poland

A total of 171 blood samples were collected from women with confirmed endometriosis, hospitalised at the Department of Surgical Gynaecology, Polish Mother’s Memorial Hospital in Lodz. A formal consent was issued by the Bioethical Committee of the Institute of the Polish Mother’s Memorial Hospital in Lodz (Approval number, 98/2015). As a control group we enrolled to the study 2934 DNA samples from POPULOUS collection of the Biobank Lab, Department of Molecular Biophysics, University of Lodz [14, 42, 43]. These samples were derived from unrelated women, who had declared themselves as a healthy, with no evidence of current endometriosis disease stage. All studied individuals were of Polish origin and provided written informed consent. The approval for their processing was obtained from The Research Bioethics Committee of University of Lodz (7/ KBBN-UL/II/2014). LODZ-Poland datasets were imputed upto 1000G P3v5.

### MELBOURNE

A total of 320 surgically confirmed cases were assessed through medical notes at the Gynaecology clinic at Royal Women’s Hospital, Melbourne, Australia. Samples were collected following written informed consent. All studies were approved by the Human Research Ethics Committee of the Royal Women’s Hospital, Melbourne, Australia (Projects 11-24 and 16-43). A control group of 887 (538 females: 349 males) controls with no history of endometriosis were selected from the population based twin study biobank from QIMR, Brisbane, Australia.

Stringent sample quality control was conducted after which, 1,207 samples remained. Samples all had high genotyping rate (>0.95), no excess heterozygosity (<3SD from mean), no first or second-degree relatives, sex checked, no outlying ancestry (all within 6 STDEVs of first 4 PCs based on European 1000 genomes phase 3 set). Followed by variant quality control, 248,472 remained after variant QC. Variants with call rate < 0.95, HWE p<10-6 and MAF < 1% were removed. Post quality control, the Melbourne dataset included 320 cases of which, 242 were stage I/II cases, 78 were stage III/IV cases and 31 were infertile cases (Infertility was defined as trying to conceive for 12 months or more on any occasion without success), and 887 controls. The genotype data was imputed using HRC Michigan imputation server up to HRC rl.l 2016.

### Northern Finland Birth Cohort (NFBC)

NFBC includes two longitudinal and prospective birth cohorts of white European women and offspring collected at 20-year intervals from the same provinces of Oulu and Lapland in Finland: NFBC1966 and NFBC1986. A total of 168 cases with a history of endometriosis were identified through national outpatient and inpatient hospital discharge registers and self-reported diagnosis through postal questionnaire at age 46 (NFBC1966). The hospital discharge registers include WHO ICD codes for identification of endometriosis disease diagnoses and dates for each hospital visit. Of the 168 cases, 55 women self-reported infertility. Controls (n=9,194) were drawn from the rest of the cohort population. Informed consent was obtained from all participants using protocols approved by the Ethical Committee of the Northern Ostrobothnia Hospital District.

The NFBC cases and controls were genotyped using the Illumina Infinium Human CNV370-Duo array. Individuals with low heterozygosity, gender-mismatch, duplicates, related individuals and SNP call rate of <0.95 or minor allele frequency (MAF) of <0.05 were excluded. The genotype data was imputed using The Haplotype Reference Consortium (HRC) reference panel, 1000 Genomes Phase 1, Michigan imputation server. Association testing was conducted in SNPTEST.

### Nurses’ Health Study II (NHS2)

NHS2 endometriosis cases were identified within the US Nurses’ Health Study II (NHS2), a prospective cohort study with ongoing follow-up from 1989. Biennially, 116,430 registered female nurses - aged 25-42 and residing in 14 of the US states at baseline - complete questionnaire information on incidence of disease outcomes and biological, environmental, dietary, and life-style risk factors. From 1996-1998, blood samples were collected from 29,613 participants 32-53 years of age. Women were asked if they had-ever had physician-diagnosed endometriosis, the date of diagnosis, and whether diagnosis had been confirmed by laparoscopy. From two validation studies (1993 and 2000) of self-reported endometriosis among these medical professional women, both demonstrated that the diagnosis was confirmed in 96% of nurses who had surgery and whose medical records were available.

The NHS2 case dataset comprised 2,230 cases with a self-reported laparoscopy-confirmed diagnosis of endometriosis and available blood samples, all of self-reported European descent. Participant enrolment, questionnaire and clinical data, and biologic sample collection have been approved by the Human Subject Committee of Harvard T.H. Chan School of Public Health and by the Institutional Review Board of Brigham and Women’s Hospital. Controls were from the Post Traumatic Stress Disorder (PTSD) GWAS within NHS2 that included 1330 women. Additionally, controls were from the Venous thromboembolism (VTE) GWAS that included a total of 4622 participants from NHS2 as well as from the Nurses’ Health Study (NHS) and the Health Professionals follow-up Study (HPFS) - prospective cohorts with data and sample collection and follow-up methods similar to NHS2. NHS is a prospective cohort of 121,700 US nurses with follow-up from 1976 when they were aged 30-45, with blood samples collected in 1989-1990 from 32,826 of these women. HPFS is a prospective cohort of 51,529 US male health professionals (e.g. physicians, dentists, pharmacists, veterinarians) aged 40 to 75 at enrollment in 1986, with blood samples collected from 18,159 men in 1993-1994.

NHS2 cases were genotyped at QIMR Berghofer Medical Research Institute, Brisbane on Illumina HumanCoreExome-12vl.O array, with ∼250,000 common tag SNPs, whereas controls were genotyped on HumanOmniExomel4vl.O (VTE) and PsychChip array (PTSD). Quality control for the GWA data resulted in the removal of SNPs with >5% missing rate, autosomal SNPs with HWE *P < 1 x* 10^−6^. Similarly, individuals with >5% missing rate, outlying heterozygosity *(+/-* three standard deviations from the mean) and of non-European ancestry based on the 1000G reference population were excluded as was one individual from each duplicate or related (pi-hat >0.2). After stringent quality control, the dataset resulted in 2,104 endometriosis cases and 5,854 controls. The dataset was imputed up to 1000G P3v5 reference panel.

### OXEGENE

A total of 1,030 cases were recruited by the Oxford Endometriosis Gene (OXEGENE) study. UK controls included 6,000 individuals from the WTCCC2. Informed consent was obtained from all participants and the study was approved by the Oxford regional multi-center and local research ethics committees.

Cases were genotyped at deCODE genetics on Illumina 670-Quad and the WTCCC2 controls were genotyped at the Wellcome Trust Sanger Institute using Illumina HumanHap 1M Beadchips. Quality control procedures for the case genotype data resulted in the removal of SNPs with genotype call rate of <0.99 and/or heterozygosity of <0.31 or >0.33. Genome-wide IBS was estimated for each pair of individuals, and one individual from each duplicate or related pair (IBS >0.82) was removed. Genotype data were combined with data from the Utah residents of Northern and Western European ancestry (CEU), Han Chinese in Beijing, China (CHB) and Japanese in Tokyo, Japan (JPT), and Yoruba from Ibadan, Nigeria (YRI) HapMap 3 reference populations, and individuals who did not have Northern European ancestry were identified using EIGENSOFT and subsequently removed. SNPs with genotype call rate of <0.95 were removed, and this threshold was increased to 0.99 for SNPs with MAF of <0.05. In addition, SNPs showing (i) deviation from HWE (*P*< 1 × 10^−6^); (ii) difference in call rate between the 1958 British Birth Cohort (58BC) and National Blood Service (NBS) control groups (*P*< 1 × 10^−4^); (iii) difference in allele and/or genotype frequency between control groups (*P*< 1 × 10^−4^); (iv) difference in call rate between cases and controls (P< 1 x 10^−4^) and (v) MAF of <0.01 were removed. Post quality control, the genotype data was pre-phased locally using SHAPEIT and the pre-phased genotype data was uploaded to the HRC Michigan imputation server for imputation to HRC rl.l 2016 reference panel. The .vcf imputed files are then used for association testing with SNPTEST where we did not include any covariates in the analysis.

### Oxford P1

Oxford P1 is composed of two centres collecting endometriosis cases and controls in the UK, namely the Oxford Endometriosis Care Centre at University of Oxford and The Women’s Centre at University of Liverpool. The ENDOX study is a prospective cohort study established at the Oxford Endometriosis CaRe Centre, University of Oxford, UK in 2012. Women undergoing diagnostic laparoscopy due to symptoms suggestive of endometriosis including pelvic pain and infertility are recruited. Endometriosis status is confirmed during surgery. The ENDOX study collects deeply phenotyped endometriosis cases and symptomatic controls using standardised World Endometriosis Research Foundation (WERF) Endometriosis Phenome and Biobanking Harmonisation Project (EPHect) guidelines [44-47], Patients complete a detailed health questionnaire on their symptomatology leading to their surgery as well as about their lifestyle and co-morbidity related medical history before attending the surgery. Before and during surgery a range of biological samples are collected. The ENDOX study was approved by National Research Ethics Service (NRES) Committee South Central Oxford (09/H0604/58b). The Liverpool study prospectively recruits women undergoing laparoscopic surgery. Cases were women who were found to have endometriosis at the surgery and patients were assigned a rAFS score based on the extended of the disease. Controls were asymptomatic women undergoing laparoscopic tubal sterilisation or having hysterectomy for uterine prolapse.

Genotyping was conducted on 194 endometriosis cases and 84 controls from the ENDOX study (177 cases : 67 controls) and the Liverpool collection (17 cases : 17 controls) using Affymetrix Axiom Human UK Biobank chip revealing 776,520 SNPs. Standard quality control (QC) was conducted [48] where samples with genotyping call-rate <0.97, outlying heterozygosity rate (3 standard deviation around mean heterozygosity rate) and ethnic outliers (non-European based clustering of samples with HapMap phase3 reference data) were excluded resulting in 165 cases (151 cases from ENDOX and 14 cases from Liverpool - 81 stage III/IV [48 infertile]; 80 stage I/II [43 infertile]; 93 infertile) and 78 controls (62 from ENDOX and 16 from Liverpool). Post-sample QC, variant QC was conducted on 773,520 SNPs in 165 cases and 78 controls. Variants with significantly different genotype call rates between cases and controls were excluded as well as variants with call rate <97%, Hardy-Weinberg equilibrium (HWE) P value of <1 x 10^−6^ or minor allele frequency (MAF) of <0.01. Furthermore, before pre-phasing, to reduce the errors due to strand misalingment, duplicates and genotyping issues with the scaffold, a stringenty QC is applied to match the scaffold against the HRC panel, removing or updating SNPs that do not agree in terms of position, alleles or frequency in the European group using the the HRC-lOOOG-check-bim.pl developed by the University of Oxford, available at http://www.well.ox.ac.uk/∼wravner/tools/. The cleaned genotype data included 655,448 SNPs was pre-phased locally using SHAPEIT v2 software and then imputed up to the HRC rl.l 2016 reference panel using the University of Michigan imputation server (https://imputationserver.sph.umich.edu/start.html). The .vcf imputed files are then used for association testing with SNPTEST where we did not include any covariates in the analysis.

### QIMRHCS

A total of 2,351 endometriosis patients were assessed from individuals recruited by The Queensland Institute of Medical Research (QIMR), Brisbane, Australia, between 1995-2002 (each completing a questionnaire and providing a blood sample). Surgical diagnosis for all endometriosis cases was confirmed from retrospective examination of medical records. Australian controls consisted of 1,870 individuals recruited by QIMR and 1,244 individuals recruited by the Hunter Community Study (HCS). Approval for the studies was obtained from the QIMR Human Ethics Research Committee, the University of Newcastle and Hunter New England Population Health Human Research Ethics Committees. Informed consent was obtained from all participants prior to testing.

QIMR cases and controls were genotyped at deCODE genetics on Illumina 670-Quad (cases) and 610-Quad (controls) Beadchips. HCS controls were genotyped at the University of Newcastle on 610-Quad Beadchips (Illumina). Genotypes for QIMR cases and controls were called with Illumina BeadStudio software. Standard quality control procedures were applied as outlined previously. Briefly, individuals with call rate of <0.95 and SNPs with mean BeadStudio GenCall score of <0.7, call rate of <0.95, Hardy-Weinberg equilibrium (HWE) *P* value of <1 x 10^−6^ or minor allele frequency (MAF) of <0.01 were excluded. Cryptic relatedness between individuals was identified through a full identity-by-state (IBS) matrix. Ancestry outliers were identified using data from 11 populations from HapMap 3 and 5 Northern European populations genotyped by the GenomeEUtwin Consortium using EIGENSOFT. To increase the power of the Australian GWAS data set, we matched the existing QIMR cases and controls by ancestry to individuals from the HCS genotyped on Illumina 610-Quad chips. After stringent quality control, the resulting QIMRHCS cohort consisted of 2,262 endometriosis cases of which 905 were stage III/IV, 1,354 stage I/II, 848 infertile and 2,924 controls. Infertility was defined as trying to conceive for 12 months or more on any occasion without success. The genotype data was imputed using HRC Michigan imputation server up to HRC rl.l 2016.

### UK Biobank

The UK Biobank (UKBB) is comprised of 500K men and women aged 40-69 at time of recruitment (2006-2010) from across the UK. The biobank was approved by the North West Multi-Centre Research Ethics Committee (MREC). In the UKBB, information was collected from participants during recruitment using questionnaires on socioeconomic status, lifestyle, family history and medical history. Participants were also followed up for cause-specific morbidity and mortality through linkage to disease registries, death registries, hospital admission records and primary care data. Also a range of biological samples including blood, urine and saliva were collected from the participants. A more detailed description of the UKBB can be found in the UK Biobank protocol (UK Biobank 2011). From this resource, we have pulled 4,196 self-report endometriosis cases of which 90% also reported a related surgery such as laparoscopy. Moreover, from medical records (ICD9/ICD10) 5027 endometriosis cases were defined (1,315 with endometriosis of ovary, intestine and rectovaginal septum - stage III/IV disease). In total, 8190 endometriosis cases of which, 1315 are stage III/IV cases and 529 who are infertile (88 self-reported and 351 with ICD9/10 codes) and all female without endometriosis or uterine fibroids, 265,265 were selected to utilised as the control set. This dataset was then restricted to individuals with genotype data and of white British ancestry bringing the final number to 6,611 endometriosis cases (1,075 stage III/IV and 417 infertile endometriosis cases) and 251,704 female controls. The UKBB genotype data passed centralised quality control and was imputed by the UKBB team up to the HRC reference panel. Given the extensive relatedness in the UKBB data, we have utilised a linear mixed model (LMM) for association testing that accounts for population structure and model the related individuals, including genotyping array as a covariate as implemented in Bolt-LMM V2.3.

### WGHS

The Women’s Genome Health Study (WGHS) is a prospective cohort of initially healthy, female North American health care professionals at least 45 years old at baseline representing participants in the Women’s Health Study (WHS) who provided a blood sample at baseline and consent for blood-based analyses. The WHS was a 2×2 trial beginning in 1992-1994 of vitamin E and low dose aspirin in prevention of cancer and cardiovascular disease with about 10 years of follow-up. Since the end of the trial, follow-up has continued in observational mode. Additional information related to health and lifestyle were collected by questionnaire throughout the WHS trial and continuing observational follow-up.

Endometriosis status was ascertained by the eighth questionnaire during the observational follow-up. WHS participants were asked: “Have you EVER had physician-diagnosed endometriosis (yes/no)? If yes, has your endometriosis diagnosis been confirmed by laparoscopy (a standard method for diagnosing endometriosis) (yes/no/unsure)?” From these questions, endometriosis status was defined two ways. A stringent definition specified cases who responded “yes” to both questions (N=613). All other endometriosis cases were defined by response “yes” to the first question, and a response of “no”, “unsure”, or “missing” to the second question (N=881). A total of 15,033 controls were defined as responding “no” to the first question. These controls were equally divided at random into two groups for pairing with the two sets of cases such that no participants were shared by analyses of two case definitions. Genotyping was conducted using the HumanHap300 Duo “+” chips; HumanHap300 Duo and iSelect chips (Illumina, San Diego, CA) with the Infinium II protocol. The genotype data was imputed using HRC Michigan imputation server up to HRC rl.l 2016.

### Genome-wide association analysis

Overall endometriosis GWAS analysis was conducted in each contributing study and subphenotype GWAS analysis was conducted for stage I/II, stage III/IV, infertile endometriosis case groups in studies with the respective case categorisation as summarised in Table 1.

### Genome-wide meta-analysis

Per-variant estimates collected from each individual study from 20,933 cases and 482,225 controls, for 10,260,082 SNPs were meta-analysed using the inverse weight variance method employed in METAL [13]. We have conducted additional three sub-phenotype meta-analyses including (1) Stage III/IV endometriosis (3,160 cases), (2) Stage I/II endometriosis (3,711 cases), (3) Infertile endometriosis (2,843 cases). Post meta-analysis, we have excluded variants that were not present in more than 50% of the effective sample size (N=8,311,853 SNPs). The genome-wide significant association p-value was determined at 5×10^−8^ and SNPs with association p<5×10^−5^ were considered suggestive associations.

### Replication analysis

The lead SNPs with p<l×10^−5^ along with two strongest proxies were selected to seek replication evidence from customers of the personal genetics company 23andMe, Inc. who had consented to participate in research. The dataset includes 37,182 cases and 251,255 controls of European ancestry. Participants were genotyped using one of four custom genotyping chips and imputed against a merged UK10K and 1000 Genomes Phase 3 reference panel.

### Gene-based association analysis

The SNP-based GWAS meta-analysis results were fed into the FUMA SNPtoGENE pipeline [15]. The gene-based association analysis utilised MAGMA software [49] for analysis, which has mapped the input SNPs to 18,742 protein-coding genes. The genome-wide significance was determined conservatively with 0.05/18742=2.67×10^−6^.

### Association of genome-wide significant Endometriosis loci with other traits/conditions Pheno Scanner

We have utilised Pheno Scanner V1.1 to identify previously established phenotype associations with genome-wide significant 27 lead SNPs from our analysis with r^2^>0.6 and p<0.001 [19]. The database was accessed on 22 June 2018.

### NHGRI GWAS Catalogue

We looked up the nearest gene to 27 genome-wide significant endometriosis SNPs for genome-wide significant association evidence with other conditions and traits using the NHGRI GWAS catalogue [20]. Database was accessed on 30 May 2018.

### Linkage Disequilibrium (LD) Score Regression

To assess the shared aetiology between endometriosis and related traits/conditions, we performed genetic correlation analysis using LD-score regression using LDHUB [23]. Publicly available genome-wide summary statistics for autoimmune, inflammatory, metabolic and pain-related diseases (N=16 ICD10 and/or self-reported conditions), anthropometric measurements (BMI, WHR), reproductive traits (Age at menarche, age at menopause, length of menstrual cycle, age of first birth, N children ever born) and pain-related symptomatology (Nine pain experiences) and seven brain regions known to be involved in pain processing were used to estimate the genome-wide genetic correlation with endometriosis. The adjusted P-value for this analysis was p<1.28×10^−3^ (0.05/39) after a Bonferroni correction.

### Biological Annotation of Associated Loci

To gather biological information on the genome-wide significant SNP, the nearest gene or genes, the cytogenetic location and summary of the gene function was collected from ANNOVAR [50]. Moreover, the nearest gene symbol was used as search term in PubMed database [51] and also in Gene Cards search tool [52] to identify relevant published literature for each of the genes that could help enlighten their role in the underlying pathophysiology of endometriosis (Accessed on 31^st^ May 2018).

### Variance explained

We estimated proportion of variance explained by a single SNP using the effect allele frequency and odds ratio from the GWA meta-analysis [32]. We used population prevalence of 8 [7, 53] and 2.5% [54] for all and stage III/IV endometriosis cases, respectively.

## Acknowledgements

We acknowledge the study participants in all the individual endometriosis studies that made this study possible. We also thank the many hospital directors and staff, gynaecologists, general practitioners and pathology services in helped with recruitment, sample collection, and confirmation of diagnoses.

We are grateful to all the families who took part in the Generation Scotland: Scottish Family Health Study, the general practitioners and Scottish School of Primary Care for their help in recruiting them, and the whole Generation Scotland team, which includes academic researchers, IT staff, laboratory technicians, statisticians and research managers. We thank staff at the University of Dundee Health Informatics Centre for their expert assistance with EHR data linkage. We thank the research participants and employees of Celmatix and 23andMe. We acknowledge the contributions from additional members of the Celmatix Research Team, including R. Mark Adams, Daniela S. Colaci, Chris Glazner, Samuel T. Globus, Sean O’Keeffe, Ursula M. Schick, Lei Tan, Cameron D. Wellock, Danielle White, and Rajeshwari R. Valiathan. Collaborators for the 23andMe Research Team are: Michelle Agee, Babak Alipanahi, Adam Auton, Robert K. Bell, Katarzyna Bryc, Sarah L. Elson, Nicholas A. Furlotte, Barry Hicks, Karen E. Huber, Ethan M. Jewett, Yunxuan Jiang, Aaron Kleinman, Keng-Han Lin, Nadia K. Litterman, Matthew H. McIntyre, Kimberly F. McManus, Joanna L. Mountain, Elizabeth S. Noblin, Carrie A.M. Northover, Steven J. Pitts, G. David Poznik, J. Fah Sathirapongsasuti, Janie F. Shelton, Suyash Shringarpure, Chao Tian, Joyce Y. Tung, Vladimir Vacic, Xin Wang, and Catherine H. Wilson.

## Supplementary Figures

**Supplementary Figure 1.** Q-Q plot for genome-wide association results for (a) Overall endometriosis, (b) stage III/IV endometriosis, (c) stage I/II endometriosis, (d) infertile endometriosis.

**Supplementary Figure 2.** Manhattan plot for genome-wide association results for endometriosis. The GWAS meta-analysis results are shown on the y-axis as -logio(P-value) and on the x-axis is the chromosomal location. The red vertical line illustrated the genome-wide significance (p<5×10^8^) and the blue vertical line shows the nominal genome-wide results (p<1×10^-5^). The genome-wide significant loci are marked with gene symbols where red are the novel results and black are previously known loci.

**Supplementary Figure 3.** Manhattan plot for genome-wide significant association results for Stage III/IV endometriosis. The GWAS meta-analysis results are shown on the y-axis as -logio(P-value) and on the x-axis is the chromosomal location. The red vertical line illustrated the genome-wide significance (p<5×10 ^8^) and the blue vertical line shows the nominal genome-wide results (p<1×10^5^).

**Supplementary Figure 4.** Manhattan plot for genome-wide significant association results for Stage I/II endometriosis. The GWAS meta-analysis results are shown on the y-axis as -logio(P-value) and on the x-axis is the chromosomal location. The red vertical line illustrated the genome-wide significance (p<5×10 ^8^) and the blue vertical line shows the nominal genome-wide results (p<1×10^-5^).

**Supplementary Figure 5.** Manhattan plot for genome-wide significant association results for infertile endometriosis. The GWAS meta-analysis results are shown on the y-axis as -logio(P-value) and on the x-axis is the chromosomal location. The red vertical line illustrated the genome-wide significance (p<5×10^8^) and the blue vertical line shows the nominal genome-wide results (p<1×10**^-5^).**

**Supplementary figure 6.** Regional association plots for novel genome-wide significant loci: (i) *DNM3* (1q24.3j, (ii) *MIIT10* (*10p12.31)* (iii) *IGF2BP3* (*7p15.3),* (iv) *RNLS* (10q23.31), (v) *GDAP1* (8q21.11), (vi) *RIN3* (14q32.12), (vii) *SKAP1* (17q21.32) for (a) endometriosis, (b) stage III/IV endometriosis, (c) stage I/II endometriosis, (d) infertile endometriosis. The association results are shown on the y-axis as -logio(P-value) and on the x-axis is the genomic location (hg 19). The top associated SNP is coloured purple and the other SNPs are coloured according to the strength of LD with the top SNP by r^2^ in the European 1000 Genomes dataset.

**Supplementary figure 7.** Regional association plots for previously established genome-wide significant loci: (i) *WNT4/CDC42* (1p36.12), (ii) *GREB1* (2p25.1), (iii) *ILIA* (2ql3), (iv) *FN1* (2q35), (v) *KDR* (4q12), (vi) *ID4* (6p22.3), (vii) *CCDC170* (6q25.1), (viii) *SYNE1* (6q25.1), (ix) *7p15.2,* (x) *7p12.3,* (xi) *CDKKN2-BAS1* (9p21.3), (xii) *FSHB* (IIp14.1), (xiii) *VEZT* (12q22) for (a) endometriosis, (b) stage III/IV endometriosis, (c) stage I/II endometriosis, (d) infertile endometriosis. The association results are shown on the y-axis as -logio(P-value) and on the x-axis is the genomic location (hg 19). The top associated SNP is coloured purple and the other SNPs are coloured according to the strength of LD with the top SNP by r^2^ in the European 1000 Genomes dataset.

**Supplementary Figure 8.** Q-Q plot for genome-wide gene-based association results for endometriosis.

**Supplementary Figure 9.** Manhattan plot for genome-wide gene-based association results for endometriosis.

**Supplementary Figure 10.** LD Score regression results for genetic enrichment between endometriosis sub-phenotypes (a) endometriosis, (b) stage III/IV endometriosis, (c) stage I/II endometriosis, (d) infertile endometriosis, and reproductive, autoimmune, metabolic and inflammatory conditions. On y-axis is the genetic correlation (rg) with standard error bars between each condition and endometriosis. On x-axis are the results per trait/condition. The red star denotes significant genetic enrichment after multiple-testing correction (p<1.28×10^3^) and green star is for nominal findings (p<0.05).

**Supplementary Figure 11.** LD Score regression results for genetic enrichment between endometriosis sub-phenotypes (a) endometriosis, (b) stage III/IV endometriosis, (c) stage I/II endometriosis, (d) infertile endometriosis, and pain symptomatology and brain volumes. On y-axis is the genetic correlation (rg) with standard error bars between each condition and endometriosis. On x-axis are the results per trait/condition. The red star denotes significant genetic enrichment after multiple-testing correction (p<1.28×10^3^) and green star is for nominal findings (p<0.05).

**Supplementary Figure 12.** LD Score regression results for genetic enrichment between endometriosis sub-phenotypes (a) endometriosis, (b) stage III/IV endometriosis, (c) stage I/II endometriosis, (d) infertile endometriosis, and reproductive traits, anthropometric traits. On y-axis is the genetic correlation (rg) with standard error bars between each condition and endometriosis. On x-axis are the results per trait/condition. The red star denotes significant genetic enrichment after multiple-testing correction (p<1.28×10^3^) and green star is for nominal findings (p<0.05).

## Supplementary Tables

**Supplementary Table 1.** Nominal genome-wide significant results (p<5×10^5^) for endometriosis.

**Supplementary Table 2.** Gene-based endometriosis association results from MAGMA-FUMA for all genes with p<0.05

**Supplementary Table 3.** Pheno Scanner look-up results for 27 genome-wide significant lead SNPs for association with other conditions/traits with r2>0.6 and p<0.001.

**Supplementary Table 4.** GWAS Catalogue look-up results for the nearest genes to genome-wide significant SNPs from Endometriosis GWAS.

